# Meta-analysis of herbicide non-target effects on pest natural enemies

**DOI:** 10.1101/2023.08.01.551535

**Authors:** Gabriel Zilnik, Paul Bergeron, Angela Chuang, Lauren Diepenbrock, Aldo Hanel, Eric Middleton, Erica Moretti, Rebecca Schmidt-Jeffris

**Affiliations:** USDA-ARS, Temperate Tree Fruit and Vegetable Crop Research Unit, 5230 Konnowac Pass Road, Wapato, WA 98951; Washington State University, 166 FSHN 100 Dairy Road, Pullman, WA 99164; University of Florida, Citrus Research and Education Center, 700 Experiment Station Rd., Lake Alfred, FL 33850; University of California, Division of Agriculture and Natural Resources, 9335 Hazard Way Suite 201, San Diego, CA 92123

**Keywords:** Herbicide, glyphosate, pesticide non-target effect, biological control, natural enemy, beneficial insects, predator, parasitoid, meta-analysis

## Abstract

A critical component of integrated pest management is minimizing disruption of biological control by reducing use of pesticides with significant non-target effects on natural enemies. Insecticide non-target effects testing for natural enemies has become increasingly common, but research examining the non-target effects of herbicides on natural enemies is scarce and recommendations regarding herbicide selectivity are non-existent. We used meta-analysis to summarize laboratory bioassays testing non-target effects of herbicides on arthropod natural enemies and identify patterns in taxon susceptibility and active ingredient toxicity. Data was extracted from 103 papers representing 801 total observations. Herbicides increased natural enemy mortality and decreased longevity, reproduction, and predation. Mesostigmatan mites and hemipterans were the most sensitive to herbicides and spiders, neuropterans, and hymenopterans were the least sensitive. Mortality was higher in juvenile predators versus parasitoids, but did not differ between adults; parasitoid juveniles are likely better protected within the host. In terms of acute mortality, metribuzin, glufosinate, and oxyfluorfen were the most harmful herbicides. Only nicosulfuron, rimsulfuron, pendimethalin, phenmedipham, atrazine, and urea did not increase natural enemy mortality. The large effect size of glufosinate is particularly concerning, as it is the most likely replacement herbicide for glyphosate in many crops. Many active ingredients remain under-studied. Our analysis indicates that herbicides have a strong potential to disrupt biological control in cropping systems.

**Simple Summary:** Reducing the use of pesticides that harm natural enemies of crop pests is important to pest management. Currently, there is limited information on how herbicides might affect natural enemies. The researchers found that herbicides increased natural enemy mortality and reduced their longevity and efficacy as predators. Some potential glyphosate replacement herbicides were more harmful than glyphosate. There was little or no data available for many herbicides and beneficial insects, indicating that much more research is needed on this topic.

## 1. Introduction

Pesticides often have non-target effects in agricultural systems [1-4]. Non-target effects include increased mortality as well as sublethal effects such as changes to behavior, development, reproduction, and molecular physiology [2]. These effects can be occur in any organism that is not the direct target of a pesticide application, including both organisms within the same taxonomic group as the target (e.g., insecticide effects on pollinators and natural enemies) or a different group (e.g., insecticide effects on vertebrate wildlife). Within the context of crop management, most of the focus of non-target effects research has focused on insecticide impacts on pollinators and natural enemies [2,5-7]. Minimizing non-target effects is paramount to developing an integrated pest management (IPM) toolkit in agricultural systems, in order to reduce disruption of biological control services provided by natural enemies [8,9]. Insecticide non-target effects research on natural enemies has increased over the decades since the introduction of the IPM conceptual framework [3,4,10].

Weed management through herbicides accounts for nearly half of all global pesticide usage [11,12]. However, unlike insecticide non-target effects, research on herbicide non-target effects on natural enemies remains scarce [1,13-16]. In the most comprehensive review of pesticide non-target effects on natural enemies, only 1.4% of records examined non-target effects of herbicides and herbicide studies did not seem to be increasing along with the other pesticide types [1]. In a recent meta-analysis of pesticide non-target effects on phytoseiids (predatory mites), only 1.6% of observations/records (*n*=2,386) and 4.5% (*n*=154) of papers examined non-target effects of herbicides [13]. In that analysis, there were an insufficient number of studies testing the same active ingredient (AI) to examine differences between herbicides [13]. A systematic review on pesticide impacts on spiders also found that relatively few herbicides had been tested [16]. This lack of research is concerning, given that there is evidence that some herbicides can be highly toxic to beneficial insects. In prior non-target effects reviews, the limited data on herbicides put these agrichemicals at or just below broad-spectrum insecticides in terms of toxicity [13,14].

Being able to compare effects of herbicide chemistries is especially important given the shift away from the use of glyphosate. Glyphosate is extensively used in both agricultural and non-agricultural settings, drawing public attention to its potential non-target effects on both human health and the environment [17-20]. In terms of beneficial arthropods, the greatest attention has been paid to the effects of glyphosate on insect pollinators, especially monarch butterflies and pollinators such as honeybees [21-24]. The impact on monarchs can be primarily attributed to the indirect effect of loss of host plants (milkweeds) on herbicide-treated agricultural systems and roadsides [21,25]. However, research on honeybees and other pollinators has a greater focus on the direct effects of glyphosate on individuals and hive health [24]. The effects of glyphosate on non-pollinator insects are not as well understood. This is especially true regarding comparisons between glyphosate and herbicides that are likely to replace it in situations where legislation, glyphosate-resistant weeds, and consumer demand cause its use to be discontinued [20,26,27]. Therefore, studies are needed that compare the effects of glyphosate and likely replacement products on beneficial insects, including pest natural enemies.

The purpose of this meta-analysis was to summarize the literature on herbicide non-target effects on a wide spectrum of arthropod natural enemies in laboratory assays. We sought to answer five research questions: (1) Do herbicides impact natural enemies through either lethal or sublethal effects, and if so, what parameters do they specifically affect? (2) Does herbicide sensitivity differ between the major natural enemy categories (predators versus parasitoids)? (3) Do taxonomic groups of natural enemies differ in herbicide sensitivity? (4) Do herbicides vary in toxicity to natural enemies based on mode of action, chemical class, or active ingredient? (5) Does glyphosate differ from likely replacement active ingredients in toxicity to natural enemies?

## 2. Materials and Methods

### 2.1. Study selection

We examined the existing literature for studies by using combinations of various keywords in GoogleScholar and Web of Science, including: herbicide, side effect, non-target, natural enemy, predator, parasitoid, insect, and arthropod. We also combined the “herbicide” search term with the scientific names of commonly tested natural enemies (e.g., “*Chrysoperla rufilabris*”, “*Trichogramma*”). Reference lists of papers retrieved in the initial search were also used as a source of additional studies. Only articles written in English or Portuguese were used; we did not find any articles written in other languages during our search. Only laboratory studies were included in our analysis; field tests of herbicide non-target effects are rare, and it is difficult to untangle the direct effect of a herbicide on natural enemies from its indirect effects (e.g., changes in habitat and food resources due to weed death) in field studies. Only formulated herbicides with named AIs were included (no experimental products or pure AI). Studies reporting proportion/percent based data were included if the percent/proportion and sample size were specified. For means-based data, studies were included if the mean, standard deviation or standard error, and sample size were specified. In some cases, only an International Organization for Biological Control rating (1-4) was provided [28]; these data could not be included. Studies only reporting only LC_50_ values were also excluded.

### 2.2. Data extraction

Data in figures was extracted using WebPlotDigitizer [29]. Herbicides were categorized into mode of action (MOA) groups and chemical classes based on the Herbicide Resistance Action Committee classifications. We simplified the response variables to 12 types: egg mortality, juvenile mortality, adult mortality, development time, longevity, size, sex ratio, mating, reproduction, predation, movement, and repellency. Only one data point for each of these response variables was used per paper, per herbicide×species combination. If more than one exposure method (e.g., contact, fresh residue, dry residue) was tested within a paper, only data from the method with the greatest risk to the natural enemy was used (contact over fresh residue, newer residues over older residues). If a variable was measured at multiple time points, only the data from the time point closest to 48 h was used, as this appeared to be the most commonly evaluated time frame (72 h data were used if both 24 and 72 h data were reported).

For mortality data, if an Abbott’s correction was used and the results for the control were not reported (common in older papers), 0% was used for the control value. Egg mortality was only used if authors assessed egg hatch directly by observing treated predator eggs. For parasitoids, any test where the infested host was treated was considered “juvenile mortality”. If more than one juvenile stage was tested in a paper, only data from the youngest stage was used; younger stages are typically the most susceptible to pesticide non-target effects due to higher surface area to volume ratios [30]. There were very few tests of herbicide exposure of pupae and these instances were not included in the analysis. The variable “Size” included measurements of weight and length or width of body parts. Sex ratio data were standardized across studies to compare the proportion of females, so that a higher effect size indicated an increase in females and a lower effect size indicated a decrease in females. Mating behavior was measured in only three studies and included ability of males to follow female cues [31] or frequency of successful mating [32,33]. The variable “Reproduction” included any study where an individual was treated and the effects on variables such as number of eggs laid, whether any offspring were produced during an observation period, number of females produced by a treated female (parasitoid), and number of adults emerging from a parasitized egg (parasitized by a treated female). “Predation” included any measure of natural enemy efficacy, including number of prey consumed or parasitized, time to first attack, and handling time. “Movement” included studies examining the effects of herbicide exposure on natural enemy speed or distance travelled. “Repellency” studies were those testing whether a natural enemy avoided a herbicide-treated surface, including amount of time spent on treated versus untreated surfaces and proportion of individuals found on treated versus untreated surfaces.

### 2.3. Data analysis

All data were analyzed using Open MEE [34]. For means data, the effect size calculated was Hedge’s *d* and for binomial data, the log odds ratio was calculated. Log odds ratios were then converted to Hedge’s *d* to allow for comparison across all studies [35]. The “Subgroup Meta-analysis” function was used to compare effect sizes between categorical variables [34]. For each categorical group analyzed, its effect size and 95% Confidence Interval (CI) were calculated. Effect sizes were considered significantly different from zero or each other when 95% CIs did not overlap.

To compare effect sizes for the different response variable types, data from all species and herbicides were combined. For the analyses comparing herbicide effects between taxonomic groups, only adult and juvenile mortality data were used (these variables had the greatest sample sizes) and all active ingredients were analyzed together. In the analyses comparing herbicide MOA group, class, and active ingredient, only entries examining adult or juvenile mortality were used and studies on all natural enemy taxa tested for each category were combined.

## 3. Results and Discussion

### 3.1. General trends in herbicide non-target research

Data from 103 papers representing 801 total observations was extracted from published literature (1960-2020) and three (at the time of analysis) unpublished studies by Schmidt-Jeffris et al. Two of the unpublished papers have since been published [36,37] and the unpublished data are noted as such in the database. No papers matching our search criteria were published prior to 1960. Of papers published from 1960-2000, only 11 papers met our search criteria, with a maximum of one publication per year. This is in contrast to 2001-2010 and 2011-2020, which had 28 and 37 suitable publications, respectively. These two decades also represented an increase in papers on the topic from approximately two per year to nearly five per year. However, publication growth was more rapid from 2001-2010 than from 2011-2020.

Overall, herbicide non-target effects research does not appear to be growing at nearly the same rate as insecticide research. In a prior meta-analysis focusing solely on phytoseiids, there was no increase in herbicide non-target effects publications throughout the years; only 1-2 papers were published in each decade where at least one herbicide was tested [13]. For insecticides, the publications typically doubled from decade-to-decade, although there was no increase this last decade [13]. Recent meta-analyses of individual insecticide classes, such as neonicotinoids and *Bt* proteins, revealed a much greater depth of research on non-target organisms than for all herbicides combined [38,39].

### 3.2. Herbicide effects on natural enemies

The most common parameters tested in studies within our database were juvenile and adult mortality (Fig. 1). Of the sublethal effects, predation, sex ratio changes, and reproduction were the most commonly examined. Across all herbicides, effect sizes for mortality of all life stages (eggs, juveniles, and adults), longevity, reproduction, and predation differed from zero, indicating that herbicides significantly change these parameters relative to a control (Fig. 1). Herbicides caused increases in mortality and decreased longevity, reproduction, and predation. Predation had a larger (more negative) effect size than reproduction (Fig. 1).

**Fig. 1.**
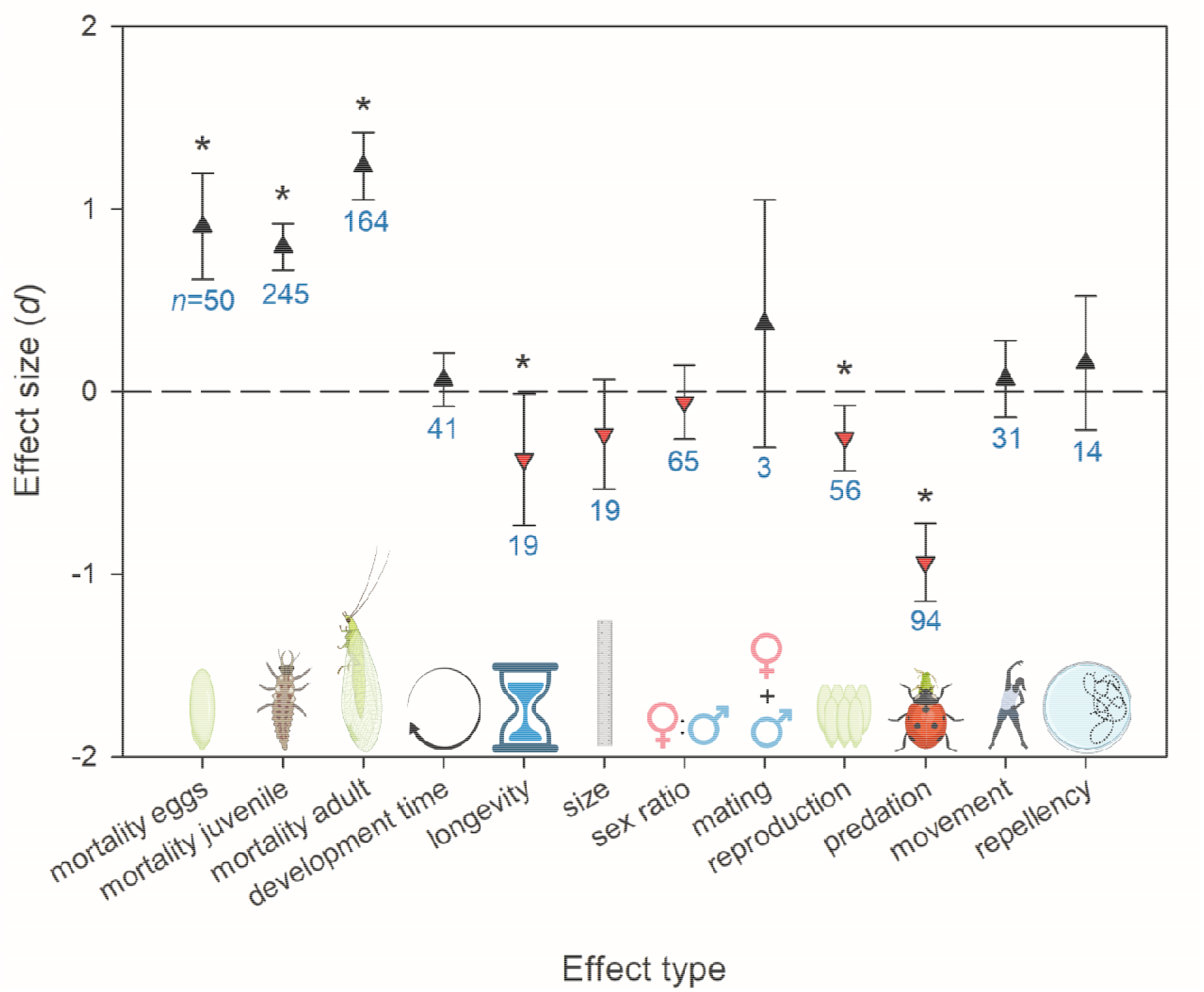
Effect sizes (*d*) for variables tested in non-target effects assays. Error bars indicate 95% bootstrapped confidence intervals. Where these intervals do not overlap with *d*=0 (dashed line), the effect is significant and indicated with an asterisk. Sample size (*n*, number of records/observations) is indicated underneath each confidence interval in blue. Positive effect sizes are indicated by black triangles pointing upwards, negative effect sizes are indicated by red triangles pointing downwards. A negative effect for sex ratio indicates a decrease in females:males compared to the control.

Previous reviews have noted that sublethal effects tend to constitute a relatively small fraction of pesticide non-target effects testing [1,2]. Within sublethal effects testing, most previous research focuses on fecundity or other measures of reproduction [1]. In our database, sublethal effects were surprisingly well represented (Fig. 1), possibly because researchers expected herbicides to typically be non-lethal at field rates and focused evaluations on sublethal effects. Predation/parasitism efficacy was the most common sublethal effect tested, with 95% of records coming from parasitoid wasps or spiders and half of records pertaining exclusively to *Trichogramma* spp. Half of all records testing predation/parasitism records were for glyphosate. While this may indicate that glyphosate is particularly disruptive to predation, it also indicates that individuals tested survived field rate applications of glyphosate long enough to have sublethal effects assessed. This is one challenge with assessing sublethal pesticide effects: highly toxic AIs do not have measurable sublethal effects at field rate because acute mortality prevents sublethal effects from being measured. Regardless, our meta-analysis shows a strong trend for herbicides to reduce the efficacy of natural enemies.

Within lethal effects, adult mortality had a larger (more positive) effect size than juvenile mortality and the egg mortality was intermediate. A previous systematic review found that adults and larvae were more sensitive than eggs and pupae [1]. This review also noted that parasitoids were the exception to this trend, where eggs and adults were more sensitive than larvae and pupae [1]. Similarly, in our meta-analysis, predators and parasitoids did not differ in adult mortality, but the juvenile mortality effect size for parasitoids was lower than that for predators (Fig. 2). This is likely because the larval stage of most parasitoids occurs within the host, where it is more protected from pesticide exposure [14]. However, the older systematic review also found that parasitoids were more pesticide susceptible than predators [1], which did not align with our results. It is possible that the trends for pesticides in general, which at the time of the review were primarily represented by broad-spectrum insecticides, do not hold for herbicides more specifically. Alternatively, there were over twice as many records of herbicide mortality effects on immature parasitoids compared to adults (Fig. 2). This has the potential to skew analysis of parasitoid taxa to indicate that they are less susceptible. To better understand how herbicides might impact parasitoids, acute effects on adults, especially of AIs other than glyphosate, should be more thoroughly investigated. This is especially true given that adults are more likely to be exposed to herbicide residues, as many adult parasitoids will forage on flowering weeds. Additionally, more research is needed on parasitoid taxa outside of Hymenoptera (e.g., dipterans) as there were no other parasitoid taxonomic groups represented in our database.

**Fig. 2.**
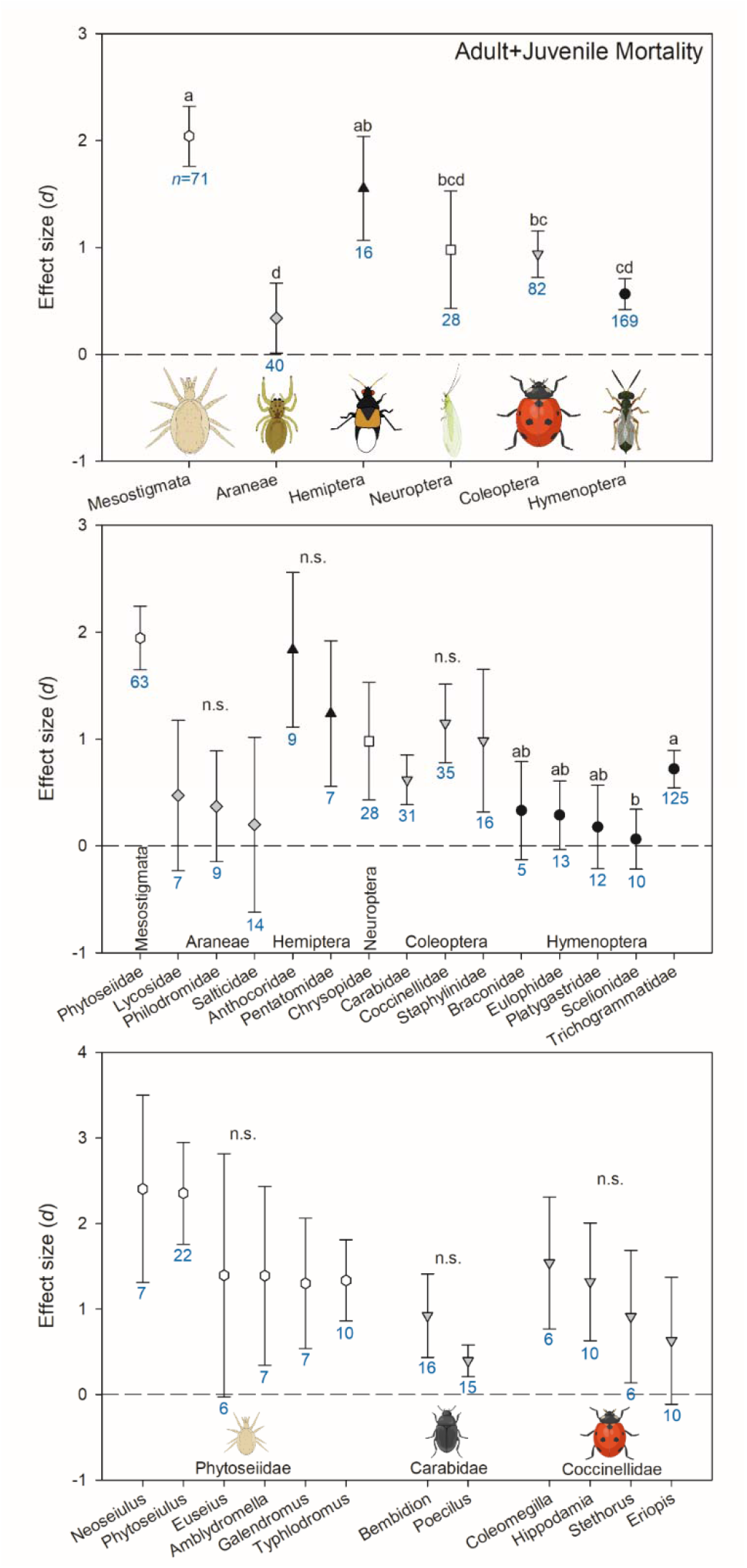
Effect sizes (*d*) of herbicide-caused mortality on both juvenile and adult stages compared between taxonomic groups at the level of (A) order, (B) families within orders, and (C) genera within families. Error bars indicate 95% bootstrapped confidence intervals. Where these intervals do not overlap with *d*=0 (dashed line), the effect is significant. Where these intervals overlap with each other, they are not considered significantly different and were assigned the same letter. Sample size (*n*, number of records/observations) is indicated underneath each confidence interva in blue. “n.s.” indicates that none of the groups compared statistically differed from each other. Symbols shade and shape for a taxonomic order is consistent between panels.

### 3.3. Taxonomic variation in herbicide effects

Hymenoptera, Coleoptera, and Mesostigmata were the best represented taxonomic orders in our database (Fig. 3A). All hymenopterans were species of parasitoid wasps and Trichogrammatidae (which were all *Trichogramma* species) was the most studied family within the order (Fig. 3B). Coccinellids, carabids, and staphylinids were the only beetle groups with published studies. Most Mesostigmata records corresponded to predatory mites in Phytoseiidae. Studies with neuropterans were limited to only chrysopids (green lacewings), with nearly all observations coming from three species of *Chrysoperla*. Within the true bugs, there was only published work on *Podisus nigrispinus* (Pentatomidae) and anthocorids (*Orius insidiosus* and *O. strigicollis*). A greater diversity of spider groups was studied, with eight different families represented in the meta-analysis. It was surprising that spiders were relatively well-represented in the herbicide non-target effects literature, given that it is often noted that spiders are under-represented in pesticide non-target effects studies more generally [1,40,41]. Otherwise, trends for which taxonomic groups of natural enemies have few or no entries in our herbicide database match the trends for other pesticides, with earwigs, non-phytoseiid predatory mites, non-coccinellid beetles, syrphids, many predatory hemipteran groups, and brown lacewings (Hemerobiidae) noticeably understudied [1]. The general lack of information on pesticide non-target effects on ground-dwelling natural enemies has been previously noted [40,42]. Because these arthropods are particularly at risk for herbicide exposure, these groups should be prioritized in future research.

**Fig. 3.**
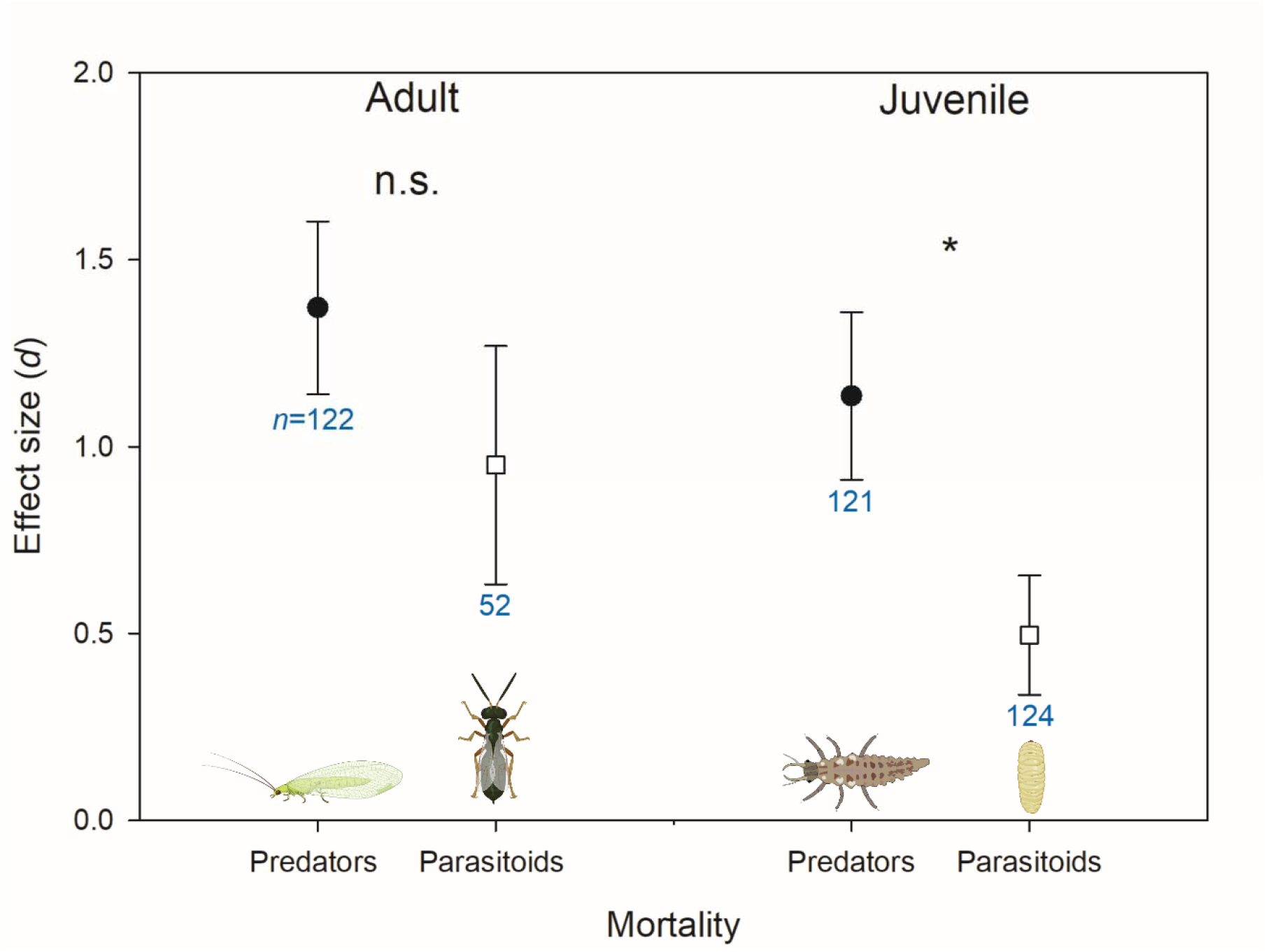
Effect sizes (*d*) of herbicide-caused mortality on adult and juvenile stages of predators versus parasitoids. Error bars indicate 95% bootstrapped confidence intervals. Where these intervals do not overlap with *d*=0, the effect is significant. Where these intervals do not overlap with each other, they are considered significantly different and marked with an asterisk. Sample size (*n*, number of records/observations) is indicated underneath each confidence interval in blue “n.s.” indicates that none of the groups compared statistically differed from each other.

Mesostigmatans and hemipterans had the highest effect sizes for combined adult and juvenile mortality, indicating higher herbicide sensitivity in these groups (Fig. 3A). Spiders and hymenopterans were the least sensitive, with lacewings and beetles intermediate. This trend also occurred at the family level; none of the spider families had an effect size that differed from zero and within the Hymenoptera, only the effect size for Trichogrammatidae differed from zero (Fig. 3B). It has been previously noted that herbicides have minimal lethal effects on spiders and most observed effects have been sublethal [16,36]. The low sensitivity of Hymenoptera may indicate that this group is particularly herbicide tolerant, or may reflect that the more protected, juvenile parasitoid stage was more commonly tested than the more sensitive adults (Fig. 2).

In an older database, mirids, ichneumonids, and chrysopids were the least susceptible to pesticides (primarily insecticides), followed by nabids, lygaeids, and anthocorids [14]. Many of these families had too few herbicide studies for analysis in our database. Of those sufficiently represented to analyze, anthocorids had the second largest effect size of any family, just slightly less than Phytoseiidae (Fig. 3B). This suggests that while anthocorids may be less sensitive to insecticides than many other groups, this trend does not necessarily extend to herbicides. This emphasizes the importance of including herbicides in non-target effects screening, as assumptions about the sensitivity of a particular group that are drawn from primarily insecticide data will not be accurate for herbicides. Our meta-analysis also supports previous qualitative reviews indicating that herbicides may be more toxic to beneficial mites than other natural enemies [43]. The authors speculate that this could be due to differing detoxification mechanisms [43], but the physiological mechanisms for herbicide non-target effects on arthropods are relatively unknown [42].

Within the orders with sufficient taxonomic represented diversity to compare families, there were few statistical differences in effect sizes (Fig. 3B). Trichogrammatids had a higher effect size than scelionids, with the other three hymenopteran families intermediate (braconids, eulophids, platygastrids) intermediate (Fig. 3B). The scelionid data were all collected in a single paper examining pesticide non-target effects on *Telenomus remus* as larvae within parasitized eggs of the host [44]. Therefore, there is not strong evidence that scelionids are less herbicide sensitive than trichogrammatids; it is possible that the herbicides tested in the single *T. remus* paper did not include as many instances of the more toxic AIs or that *T. remus* may be unusually herbicide tolerant compared to scelionids in general. Similar to the analysis within orders, within taxonomic families there were no differences in effect size for combined adult and juvenile mortality between genera, although only three families (Phytoseiidae, Carabidae, and Coccinellidae) had multiple genera sufficiently represented in the dataset to allow for comparisons (Fig. 3C). Therefore, based on currently available data, it appears that significant differences in herbicide susceptibility occur almost entirely at the order level. This may indicate that generalizations about toxicity of particular herbicides can be made at the order level. If so, future studies should prioritize testing herbicide effects on understudied orders of natural enemies [42] or testing the effects of understudied AIs, before testing additional species or families in well-studied groups.

### 3.4. Variation between herbicides

Herbicide MOA groups differed in toxicity to natural enemies. Group 9 (inhibitors of EPSP synthase) made up a large proportion of the data set (27%) due to an overrepresentation of studies on glyphosate (Fig. 4A). Group 10 (inhibitors of glutamine synthetase) was the most toxic to natural enemies and Groups 3 (microtubule assembly inhibitors) and 27 (inhibitors of 4-HPPD) were the least toxic; these were also the only two groups with effect sizes that did not differ from zero (Fig. 4A).

**Fig. 4.**
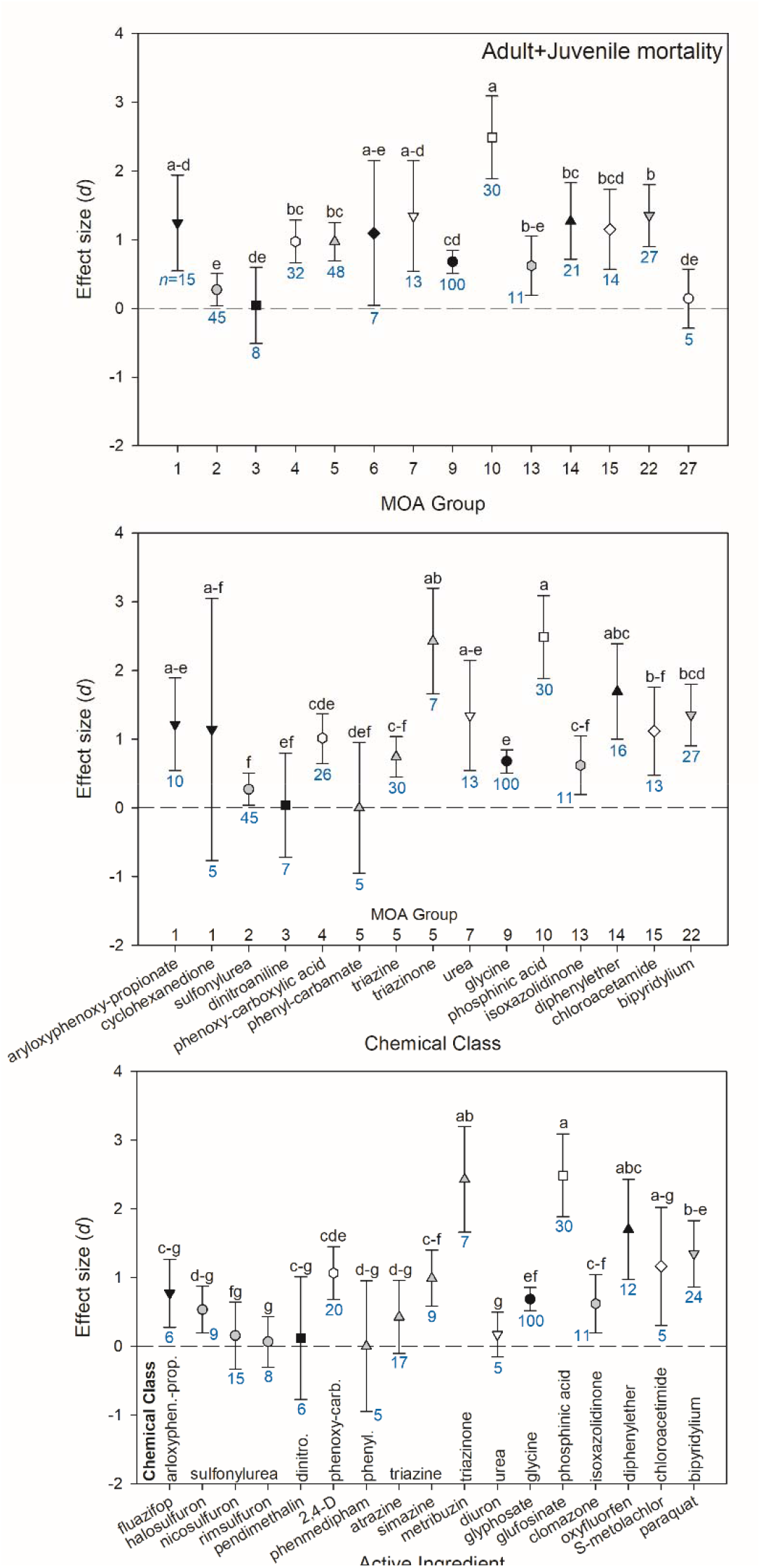
Effect sizes (*d*) of herbicide-caused mortality on both juvenile and adult stages compared between herbicide (A) mode of action (MOA) group, (B) chemical class, and (C) active ingredient. Error bars indicate 95% bootstrapped confidence intervals. Where these intervals do not overlap with *d*=0 (dashed line), the effect is significant. Where these intervals overlap with each other, they are not considered significantly different and were assigned the same letter. Sample size (*n*, number of records/observations) is indicated underneath each confidence interval in blue. “n.s.” indicates that none of the groups compared statistically differed from each other. Symbols shade and shape for a MOA group is consistent between panels.

Only two MOA groups had more than one chemical class adequately represented in the data set for within-group comparisons: Groups 1 (inhibitors of acetyl CoA carboxylase) and 5 (inhibitors of photosynthesis at photosystem II). Within Group 1, aryloxyphenoxy-propionates (FOPs) and cyclohexanediones (DIMs) did not differ in effect size, although the effect size variance for cyclohexanediones was very large (Fig. 4B). Within Group 5, the triazinones had a higher effect size that the phenyl-carbamates and triazines. Across all chemical classes, sulfonylureas, dinitroanilines, phenyl-carbamates, triazines, glycines, isoxazolidinones, and chloroacetamides had the lowest effect sizes (Fig. 4B). Conversely, phosphinic acids, triazinones, and diphenylethers had the highest effect sizes (Fig. 4B). Some chemical classes were only represented by one active ingredient, such as phenyl-carbamates (phenmedipham), triazinones (metribuzin), glycines (glyphosate), and isoxazolidinone (clomazone).

For two chemical classes in the data set, there were enough observations to compare active ingredients within a class for combined adult and juvenile mortality. The sulfonylureas (halosulfuron, nicosulfuron, and rimsulfuron) did not differ from each other in effect size (Fig. 4C). The triazines, atrazine and simazine, also did not differ in effect size (Fig. 4C). Across all active ingredients, the herbicides least acutely toxic to natural enemies (effect size with confidence interval overlapping zero) were nicosulfuron, rimsulfuron, pendimethalin, phenmedipham, atrazine, and diuron. The effect size for glyphosate was relatively low, although it did differ from zero. Metribuzin and glufosinate had the largest effect sizes, and oxyfluorfen and S-metolachlor did not statistically differ from those active ingredients, although S-metolachlor had a fairly high effect size variance and low sample size (Fig. 4C). Paraquat had a significantly lower effect size than glufosinate but did not differ from the other three most harmful active ingredients. Therefore, our meta-analysis indicates that glufosinate, metribuzin, oxyfluorfen, and paraquat are the most harmful herbicide AIs to natural enemies and additional work is needed to determine whether S-metolachlor should also be considered part of this group.

There has been little prior work attempting to discern patterns regarding which herbicides are most harmful to natural enemies. In the older systematic review, the nitrophenol derivative, nitrogen heterocyclic, urea derivative, and carbamate chemical classes were fairly toxic, whereas organometallics and phenoxy-alkyl derivates were not toxic [14]. A qualitative summary of herbicide non-target effects on phytoseiids found that fomofenoxim, bromoxynil, ioxynil, methabenzthiazuron, glufosinate, and paraquat were consistently harmful [45]. Based on our analysis, herbicide toxicity appears to be primarily driven by chemical class; chemical classes within a single MOA differed in effect size (MOA 5), whereas active ingredients within the same chemical class did not differ in effect size (sulfonylureas, triazines). This is not unexpected, as MOA groups for herbicides are based on mechanism of toxicity to plants, not animals.

Therefore, effects of herbicides on arthropods may be generalizable at the chemical class level. However, given the relatively few groups for which there were adequate records to perform this analysis, more work will be needed to confirm this generalization.

“Inert ingredients” within formulated pesticides further complicate identifying patterns within herbicide toxicity. It is important that non-target effects researchers test formulated products, as natural enemies will never be exposed to pure AI in the field. However, research examining pesticide non-target effects on bees has found that differences between formulations with the same AI can alter conclusions of non-target effects testing [46]. Surfactants and other adjuvants, either included in the pesticide formulation, or tank mixed to improve herbicide performance can increase AI toxicity or even have direct, non-synergistic impacts on natural enemies [37,45,47,48]. In the case of glyphosate, much of its reported toxicity to bees is attributed to the inert, surfactant ingredient ethoxylated tallowamine [46]. While our meta-analysis focuses on formulated herbicides, there is a clear need for comparing different herbicide formulations, inert ingredients, and pure AI in order to determine mechanisms of herbicide toxicity. This may allow for the modification of existing herbicide formulations to reduce harm to natural enemies and other non-target organisms.

Toxicity of individual AIs may partially drive patterns observed in taxonomic group susceptibility. In the combined juvenile + adult mortality analysis, 37% of the Hymenoptera records tested glyphosate. This likely reflects the intersection of high glyphosate use in field crops and the importance of *Trichogramma* spp. for caterpillar biological control within these cropping systems, particularly in Brazil, where much of this research was conducted [49]. For the other arthropod orders, glyphosate comprised only 11-21% of records. Similarly, within the juvenile + adult mortality analysis, Hymenoptera had relatively fewer records (13%) pertaining to the four most harmful AIs (glufosinate, metribuzin, oxyfluorfen, paraquat) compared to most other taxonomic orders (25-31% of records, with the exception of Coleoptera which also had 13%). Therefore, an over-representation of the less harmful glyphosate and an under-representation of the four most harmful AIs, combined with an emphasis on testing the juvenile (more protected) stage may make Hymenoptera appear less herbicide sensitive than other groups in our database.

### 3.5. Glyphosate alternatives

Glyphosate is the most widely used herbicide [19] primarily due to the adoption of glyphosate-resistant row crops and the availability of low cost, generic formulations [20]. It is therefore unsurprising that 32% of our records tested non-target effects of this AI. Globally, there is widespread interest in reducing glyphosate use due weed resistance development, and more recently, concerns regarding environmental safety and human health [20,26,27].

Weed management alternatives to glyphosate rely on multiple herbicides and are often still less effective than the glyphosate applications they replace [27]. Glyphosate is also less acutely toxic to mammals than most other herbicides [50]. Paraquat is the main burn-down alternative to glyphosate in field crops [20], although it is less effective [19] and is also one of the most toxic herbicides to humans and other vertebrates [51]. In tree fruit, paraquat and glufosinate are the most likely glyphosate replacements for managing intra- and inter-row weeds post-emergence [37]. In the European Union (EU), only quizalofop-ethyl is available as a post-emergent herbicide alternative to glyphosate in tree fruit and cycloxydim, 2,4-D, and dicamba are the most suitable replacements in herbaceous crops [27]. However, the latter three herbicides must be used in combination as cycloxydim is only effective against monocots and 2,4-D and dicamba are only effective against dicots [27]. Field crop varieties resistant to glufosinate, dicamba, 2,4-D, triazines, and imidazolinones are also available and increased used of these varieties and their respective herbicides is expected if glyphosate is banned in some countries [19,20].

Based on the meta-analysis results, replacement of glyphosate with other herbicide AIs would likely be disruptive to biological control, especially outside of the EU. Glufosinate and paraquat were among the most acutely AIs to natural enemies. The triazine class (Fig. 4B) and 2,4-D (Fig. 4C) also significantly increased mortality of natural enemies. There were no studies in our database that tested effects of imidazolinone herbicides. Quizalofop-ethyl was classified as harmless to two predatory beetles [52], but there have been no additional studies on this AI. Cycloxidim was also poorly represented in our database; exposure caused high mortality in adult *Trichogramma cacoeciae* [53], but no mortality on *Bembidion lampros* [52]. Surprisingly, there are also very few studies on the effects of dicamba. Previous research has found mixed results; exposure caused no adult mortality in *T. cacoeciae* [53], increased male mortality in *Coleomegilla maculata* [54], and increased female mortality in *Phytoseiulus persimilis* [45]. It has been suggested that lethal effects of dicamba are primarily caused by inert ingredients, leading formulations to differ in their toxicity [54]. Therefore, likely glyphosate replacement herbicides appear to be highly toxic to natural enemies (especially the primary burndown AIs, glufosinate and paraquat) or are very poorly studied. Non-target effects tests of paraquat exposure in predatory mites remains one of the only field studies linking herbicide use to secondary pest outbreaks [55,56]. Field tests of other likely glyphosate replacement chemistries, and non-chemical management strategies, are needed to determine which weed management programs are the least disruptive to biological control.

## 4. Conclusion

There is movement towards whole-systems IPM to better understand how other aspects of crop management can affect the management of arthropod pests. This includes aspects of weed management like tillage and mowing, but often neglects herbicides. No-tillage and reduced tillage systems frequently increase beneficial insects and reduce the need for chemical insect control [57-59]. However, reduced plant diversity due to strict weed management could lead to fewer generalist predators which may harm biological control services [60]. Translating non-target bioassay results from the laboratory to the field is always challenging [42], but herbicides may pose a unique challenge. A decrease in natural enemy abundance following an herbicide application may be due to toxicity, but it could also be caused by loss of habitat or other resources provided by treated weeds that results in natural enemy dispersal. Weed management has been shown to have an impact on arthropod populations including both pest and beneficial insects [61,62]. Herbicides may be considered the least selective of all pesticides because they “directly affect primary producers, with subsequent effects rippling through dependent trophic levels in the community” [63].

## Supplementary Materials

The following supporting information can be downloaded at: www.mdpi.com/xxx/s1**, Database S1:** Herbicide non-target effects database used to conduct the meta-analysis.

## Author Contributions

Conceptualization, R.S.; methodology, R.S.; formal analysis, R.S.; investigation, G.Z., P.B., A.C., L.D., A.H., E.M., E.M., R.S.; data curation, R.S..; writing— original draft preparation, R.S. and G.Z.; writing—review and editing, G.Z., P.B., A.C., L.D., A.H., E.M., E.M., R.S.; visualization, R.S.; supervision, R.S. and L.D.; project administration, R.S. All authors have read and agreed to the published version of the manuscript.

## Data Availability Statement

The data supporting these results is provided in the supplementary material “Database S1”.

## Acknowledgements

We thank C. Sater with assistance on data extraction from papers. Figures created with BioRender.com. The use of trade, firm, or corporation names in this publication is for the information and convenience of the reader. Such use does not constitute an official endorsement or approval by the United States Department of Agriculture or the Agricultural Research Service of any product or service to the exclusion of others that may be suitable.

## References

1. Theiling, K.M.; Croft, B.A. Pesticide side-effects on arthropod natural enemies: a database summary. Agric. Ecosyst. Environ. 1988, 21, 191–218.

2. Desneux, N.; Decourtye, A.; Delpuech, J.-M. The sublethal effects of pesticides on beneficial arthropods. Annu. Rev. Entomol. 2007, 52, 81–106.

3. Croft, B.A. Natural enemies and pesticides: an overview. In Arthropod Biological Control Agents and Pesticides; John Wiley & Sons, Inc.: New York, NY, 1990; pp. 3–15.

4. Pimentel, D. Ecological effects of pesticides on non-target species. Executive office of the President, Office of Science and Technology, Washington, D.C.; 1971.

5. Brittain, C.; Potts, S.G. The potential impacts of insecticides on the life-history traits of bees and the consequences for pollination Basic and Applied Ecology 2011, 321-331.

6. Godfray, H.C.J.; Blacquiere, T.; Field, L.M.; Hails, R.S.; Petrokofsky, G.; Potts, S.G.; Raine, N.E.; Vanbergen, A.J.; McLean, A.R. A restatement of the natural science evidence base concerning neonicotinoid insecticides and insect pollinators. Proc. R. Soc. B. 2014, 281, 20140558.

7. Egan, P.A.; Dicks, L.V.; Hokkanen, H.M.; Stenberg, J.A. Delivering integrated pest and pollinator management (IPPM). Trends in Plant Science 2020, 25, 577–589.

8. Hull, L.A.; Beers, E.H. Ecological selectivity: modifying chemical control practices to preserve natural enemies. In Biological Control in Agricultural IPM Systems, Hoy, M.A., Herzog, D.C., Eds.; Academic Press: New York, NY, 1985; pp. 103–122.

9. Torres, J.B.; Bueno A.d.F. Conservation biological control using selective insecticides – A valuable tool for IPM. Biol. Control 2018, 126, 53–64, doi:10.1016/j.biocontrol.2018.07.012.

10. Doutt, R.L. Ecological considerations in chemical control: implications to non-target invertebrates. Bull. Entomol. Soc. Am. 1964, 10, 83–88.

11. Sharma, A.; Kumar, V.; Shahzad, B.; Tanveer, M.; Sidhu, G.P.S.; Handa, N.; Kohli, S.K.; Yadav, P.; Bali, A.S.; Parihar, R.D.;, et al. Worldwide pesticide usage and its impacts on ecosystem. SN Appl. Sci. 2019, 1, 1446.

12. FAO. License: CC BY-NC-SA 2.0 IGO. Extracted from https://www.fao.org/faostat/. Date of Access: 02-15-2022. 2022.

13. Schmidt-Jeffris, R.A.; Beers, E.H.; Sater, C. Meta-analysis and review of pesticide non-target effects on phytoseiids, key biological control agents. Pest. Manag. Sci. 2021, 77, 4848–4862, doi:10.1002/ps.6531.

14. Croft, B.A. Pesticide effects on arthropod natural enemies: a database summary. In Arthropod Biological Control Agents and Pesticides; John Wiley & Sons, Inc.: New York, NY, 1990; pp. 17–46.

15. Fountain, M.T.; Medd, N. Integrating pesticides and predatory mites in soft fruit crops. Phytoparasitica 2015, 43, 657–667.

16. Pekar, S. Spiders (Araneae) in the pesticide world: an ecotoxicological review. Pest Manag. Sci. 2012, 68, 1438–1446, doi:10.1002/ps.3397.

17. Benbrook, C.M. Trends in glyphosate herbicide use in the United States and globally. Environmental Sciences Europe 2016, 28, 1–15.

18. Myers, J.P.; Antoniou, M.N.; Blumberg, B.; Carroll, L.; Colborn, T.; Everett, L.G.; Hansen, M.; Landrigan, P.J.; Lanphear, B.P.; Mesnage, R.;, et al. Concerns over use of glyphosate-based herbicides and risks associated with exposures: a consensus statement. Environmental Health 2016, 15, 1–13.

19. Duke, S.O. The history and current status of glyphosate. Pest Manag. Sci. 2018, 74, 1027–1034.

20. Beckie, H.J.; Ken C. Flower; Ashworth., M.B. Farming without glyphosate? Plants 2020, 9, 96.

21. Pleasants, J.M.; Oberhauser, K.S. Milkweed loss in agricultural fields because of herbicide use: effect on the monarch butterfly population. Insect Conservation and Diversity 2013, 6, 135–144.

22. Balbuena, M.S.; Tison, L.; Hahn, M.-L.; Greggers, U.; Menzel, R.; Farina, W.M. Effects of sublethal doses of glyphosate on honeybee navigation. J. Exp. Biol. 2015, 218, 2799–2805.

23. Abraham, J.; Benhotons, G.S.; Krampah, I.; Tagba, J.; Amissah, C.; Abraham, J.D. Commercially formulated glyphosate can kill non-target pollinator bees under laboratory conditions. Entomol. Exp. Appl. 2018, 166, 695–702.

24. Battisti, L.; Potrich, M.; Sampaio, A.R.; de Castilhos Ghisi, N.; Costa-Maia, F.M.; Abati, R.; Dos Reis Martinez, C.B.; Sofia, S.H. Is glyphosate toxic to bees? A meta-analytical review. Sci Total Environ 2021, 767, 145397, doi:10.1016/j.scitotenv.2021.145397.

25. Stenoien, C.; Nail, K.R.; Zalucki, J.M.; Parry, H.; Oberhauser, K.S.; Zalucki., M.P. Monarchs in decline: a collateral landscape-level effect of modern agriculture. Insect Sci 2018, 25, 528–541.

26. Beckie, H.J. Herbicide-resistant weed management: focus on glyphosate. Pest Manag. Sci. 2011, 67, 1037–1048.

27. Fogliatto, S.; Ferrero, A.; Vidotto, F. Current and future scenarios of glyphosate use in Europe: Are there alternatives? Advances in Agronomy 2020, 163, 219–278.

28. Jansen, J.-P. Pest Select Database: a new tool to use selective pesticides for IPM. Comm. Appl. Biol. Sci. 2013, 78, 115–119.

29. Rohatgi, A. WebPlotDigitizer 4.2. Available from: https://automeris.io/WebPlotDigitizer. 2019.

30. Croft, B.A. Factors affecting susceptibility. In Arthropod Biological Control Agents and Pesticides; John Wiley & Sons, Inc.: New York, NY, 1990; pp. 71–100.

31. Leccia, F.; Kysilková, K.; Kolářová, M.; Hamouzová, K.; Líznarová, E.; Korenko, S. Disruption of the chemical communication of the European agrobiont ground-dwelling spider *Pardosa agrestis* by pesticides. J. Appl. Entomol. 2016, 140, 609–616, doi:10.1111/jen.12288.

32. Michalková, V.; Pekár, S. How glyphosate altered the behaviour of agrobiont spiders (Araneae: Lycosidae) and beetles (Coleoptera: Carabidae). Biol. Control 2009, 51, 444–449, doi:10.1016/j.biocontrol.2009.08.003.

33. Godfrey, J.A.; Rypstra, A.L. Impact of an atrazine-based herbicide on an agrobiont wolf spider. Chemosphere 2018, 201, 459–465, doi:10.1016/j.chemosphere.2018.03.023.

34. Wallace, B.C.; Lajeunesse, M.J.; Dietz, G.; Dahabreh, I.J.; Trikalinos, T.A.; Schmid, C.H.; Gurevitch, J.; Poisot, T. Open MEE: Intuitive, open-source software for meta-analysis in ecology and evolutionary biology. Methods Ecol. Evol. 2017, 8, 941–947, doi:10.1111/2041-210x.12708.

35. Borenstein, M.; Hedges, L.V.; Higgins, J.P.T.; Rothstein, H.R. Converting among effect sizes. In Introduction to Meta-Analysis; John Wiley & Sons, Ltd: West Sussex, UK, 2009; pp. 45–49.

36. Schmidt-Jeffris, R.A.; Moretti, E.A.; Bergeron, P.E.; Zilnik, G. Nontarget impacts of herbicides on spiders in orchards. J Econ Entomol 2022, 115, 65–73, doi:10.1093/jee/toab228.

37. Bergeron, P.; Schmidt-Jeffris, R. Herbicides harm key orchard predatory mites. Insects 2023, 14, doi:10.3390/insects14050480.

38. Main, A.R.; Webb, E.B.; Goyne, K.W.; Mengel, D. Neonicotinoid insecticides negatively affect performance measures of non-target terrestrial arthropods: A meta-analysis.Ecol. Appl. 2018, 28, 1232–1244.

39. Naranjo, S.E. Effects of GE crops on non-target organisms. Plant biotechnology: Experience and future prospects 2021, 127–144.

40. Overton, K.; Hoffmann, A.A.; Reynolds, O.L.; Umina, P.A. Toxicity of insecticides and miticides to natural enemies in Australian grains: A review. Insects 2021, 12, 187, doi:10.3390/insects12020187.

41. Pekar, S.; Haddad, C.R. Can agrobiont spiders (Araneae) avoid a surface with pesticide residues? Pest Manag Sci 2005, 61, 1179–1185, doi:10.1002/ps.1110.

42. Schmidt-Jeffris, R.A. Nontarget pesticide impacts on pest natural enemies: progress and gaps in current knowledge. Curr Opin Insect Sci 2023, 58, 101056, doi:10.1016/j.cois.2023.101056.

43. Gerson, U.; Smiley, R.L.; Ochoa, R. The effect of agricultural chemicals on acarine biocontrol agents. In Mites (Acari) for Pest Control; Blackwell Science: Oxford, UK, 2003; pp. 367–383.

44. Carmo, E.L.; Bueno, A.F.; Bueno, R.C.O.F. Pesticide selectivity for the insect egg parasitoid *Telenomus remus*. BioControl 2010, 55, 455–464, doi:10.1007/s10526-010-9269-y.

45. Schmidt-Jeffris, R.A.; Cutulle, M.A. Non-target effects of herbicides on *Tetranychus urticae* and its predator, *Phytoseiulus persimilis*: implications for biological control. Pest Manag. Sci. 2019, 75, 3226–3234, doi:10.1002/ps.5443.

46. Mullin, C.A. Effects of ‘inactive’ ingredients on bees. Curr Opin Insect Sci 2015, 10, 194–200, doi:10.1016/j.cois.2015.05.006.

47. Niedobová, J.; Skalský, M.; Ouředníčková, J.; Michalko, R. Glyphosate based formulation with tank mixing adjuvants alter predatory behaviour of the ground dwelling lycosid spider *Pardosa* sp. IOBC/WPRS Bulletin 2019, 143, 20–24.

48. Niedobova, J.; Skalsky, M.; Ourednickova, J.; Michalko, R.; Bartoskova, A. Synergistic effects of glyphosate formulation herbicide and tank-mixing adjuvants on *Pardosa* spiders. Environ Pollut 2019, 249, 338–344, doi:10.1016/j.envpol.2019.03.031.

49. Rakes, M.; Pasini, R.A.; Morais, M.C.; Araujo, M.B.; de Bastos Pazini, J.; Seidel, E.J.; Bernardi, D.; Grutzmacher, A.D. Pesticide selectivity to the parasitoid *Trichogramma pretiosum*: A pattern 10-year database and its implications for Integrated Pest Management. Ecotoxicol. Environ. Saf. 2021, 208, 111504, doi:10.1016/j.ecoenv.2020.111504.

50. Kniss, A.R. Long-term trends in the intensity and relative toxicity of herbicide use. Nat Commun 2017, 8, 14865, doi:10.1038/ncomms14865.

51. Summers, L.A. The Bipyridinum Herbicides; Academic Press: London, UK, 1980.

52. Jansen, J.-P.; Hautier, L.; Mabon, N.; Schiffers, B. Pesticides selectivity list to beneficial arthropods in four field vegetable crops. IOBC WPRS Bulletin 2008, 35, 66–77.

53. Hassan, S.A.; Hafes, B.; Degrande, P.E.; Herai, K. The side-effects of pesticides on the egg parasitoid *Trichogramma cacoeciae* Marchal (Hym., Trichogrammatidae), acute dose-response and persistence tests. J. Appl. Entomol. 1998, 122, 569–573, doi:10.1111/j.1439-0418.1998.tb01547.x.

54. Freydier, L.; Lundgren, J.G. Unintended effects of the herbicides 2,4-D and dicamba on lady beetles. Ecotoxicology 2016, 25, 1270–1277, doi:10.1007/s10646-016-1680-4.

55. Metzger, J.A.; Pfeiffer, D.G. Topical toxicity of pesticides used in Virginia vineyards to the predatory mite, *Neoseiulus fallacis* (Garman). J. Entomol. Sci. 2002, 37, 329–337.

56. Pfeiffer, D.G. Effects of field applications of paraquat on densities of *Panonychus ulmi* (Koch) and *Neoseiulus fallacis* (Garman). J. Agric. Entomol. 1986, 3, 322–325.

57. House, G.J.; Stinner, B.R. Arthropods in no-tillage soybean agroecosystems: community composition and ecosystem interactions. Environmental Management 1983, 7, 23–28.

58. Rowen, E.; Regan, K.; Barbercheck, M.; Tooker, J.F. Is tillage beneficial or detrimental for invertebrate pest management? A meta-analysis. *Agriculture*, Ecosystems, and Environment 2020, 294, 106849.

59. House, G.J., and G. E. Brust. Ecology of low-input, no-tillage agroecosystems. Agric. Ecosyst. Environ. 1989, 27, 331–345.

60. Dassou, A.G.; Tixier, P. Response of pest control by generalist predators to local-scale plant diversity: a meta-analysis. Ecol Evol 2016, 6, 1143–1153, doi:10.1002/ece3.1917.

61. BÀRberi, P.; Burgio, G.; Dinelli, G.; Moonen, A.C.; Otto, S.; Vazzana, C.; Zanin, G. Functional biodiversity in the agricultural landscape: relationships between weeds and arthropod fauna. Weed Res. 2010, 50, 388–401, doi:10.1111/j.1365-3180.2010.00798.x.

62. Norris, R.F.; Kogan, M. Ecology of interactions between weeds and arthropods. Annu Rev Entomol 2005, 50, 479–503, doi:10.1146/annurev.ento.49.061802.123218.

63. Croft, B.A. Ecological influences. In Arthropod Biological Control Agents and Pesticides; John Wiley & Sons: New York, NY, 1990; pp. 185–218.

